# Next generation digital PCR: high dynamic range single molecule DNA counting via ultra-partitioning

**DOI:** 10.1101/2022.08.01.502071

**Authors:** Eleen Y. Shum, Janice H. Lai, Sixing Li, Haeun G. Lee, Jesse Soliman, Vedant K. Raol, Cavina K. Lee, Stephen P.A. Fodor, H. Christina Fan

**Affiliations:** Enumerix, Inc. 4030 Fabian Way, Palo Alto, CA 94070

**Keywords:** Single molecule, gene expression, DNA counting, digital PCR, lightsheet microscopy, 3D imaging, emulsion, NIPT, fetal aneuploidy

## Abstract

Digital PCR (dPCR) was first conceived for single molecule quantitation. However, current dPCR systems often require DNA templates to share partitions due to limited partitioning capacities. Here, we introduce Ultra-dPCR, a next generation dPCR system where DNA counting is performed in a single-molecule regimen through a 6-log dynamic range using a swift and parallelized workflow. Each Ultra-dPCR reaction is divided into >30 million partitions without microfluidics to achieve single template occupancy. Combined with a unique emulsion chemistry, partitions are optically clear and enable the use of a 3D imaging technique to rapidly detect DNA-positive partitions. Single molecule occupancy also allows for more straightforward multiplex assay development due to the absence of partition-specific competition. As a proof-of-concept, we developed a 222-plex Ultra-dPCR assay and demonstrated its potential use as a rapid, low-cost screening assay for non-invasive prenatal testing (NIPT) for as low as 4% trisomy fraction samples with high precision, accuracy, and reproducibility.

---

One of the most transformative methods for precise DNA quantification is digital PCR (dPCR), which originates from the concept of isolating single DNA molecules into individual compartments for amplification and detection^1–5^. In the early studies, DNA samples were diluted and quantified using PCR in a limited dilution protocol, which involved a highly laborious partitioning process^5,6^. With advances in microfluidics, modern dPCR platforms have emerged that take advantage of various methods for DNA partitioning ^7–9^. Partitioning-based counting not only simplifies the way to count DNA, but also improves the sensitivity of rare molecule detection and PCR signal amplitude even in the presence of PCR inhibitors^10^.

Despite these advancements, current dPCR systems offer relatively low partition capacity (≤ 100K) per sample, due to physical limitations and manufacturing constraints^11,12^. In many applications, DNA quantitation with these systems often occurs outside of the single molecule domain, where >1 target molecules occupy the same partition (Figure 1). In this regime, multiple adaptations are required for a successful dPCR experiment. First, Poisson statistics is used to estimate the original DNA input. This necessitates the determination of both DNA-positive and DNA-negative partitions ^11–13^, which can increase workflow time per sample and/or limit partition imaging capacity. Poisson correction has its limitations especially in high DNA occupancy regimes, as quantitative precision reduces exponentially until saturation of all partitions is achieved^12^. Second, when multiple DNA templates are housed in the same partitions, partition-specific competition (PSC) can occur in multiplexing assays, where two or more amplicons compete for limited PCR reagents^12,13^. To reduce the effect of PSC, multiplex dPCR assays require a more rigorous assay optimization process to balance multiplex PCR efficiencies^13^. Altogether, current dPCR systems have found success in applications where the DNA dynamic range requirement is low, and assays can be standardized, such as in clinical workflows for cancer liquid biopsy and infectious diseases^14^.

**Figure 1.**
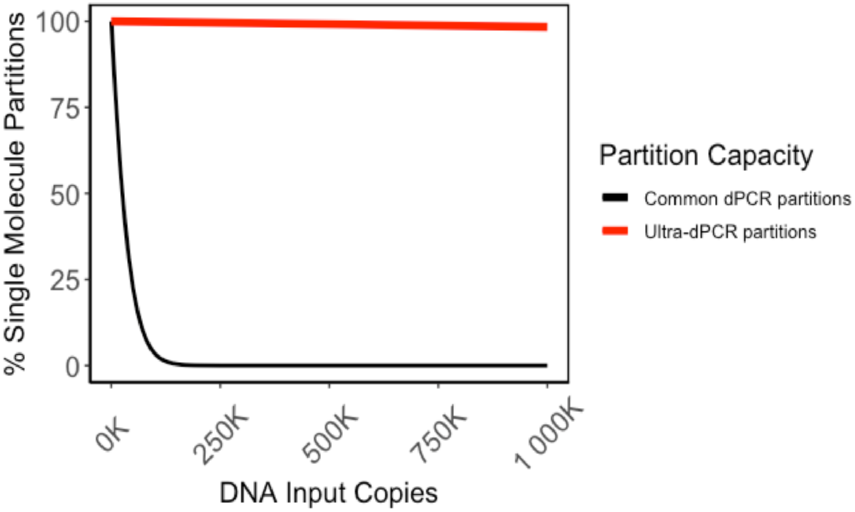
Single molecule partition vs DNA input. Partitions with a single DNA molecule over all DNA-positive partitions are calculated based on Poisson statistics for a common partition capacity of 20K partitions^11^ versus Ultra-dPCR’s 30 million partitions. Low partition systems fall out of the single molecule domain rapidly and require Poisson correction to derive quantitative accuracy up to a threshold number of counts^21,22^. In contrast, even at 1 million DNA input copies, Ultra-dPCR still maintains the limited dilution regime, allowing for true single molecule counting.

The ideal dPCR system would involve a significantly greater number of partitions to reduce or eliminate partition saturation, thereby increasing the quantitative precision and dynamic range^12^. Many high partition dPCR systems have had limited commercial success, due to prohibitively high cost, low throughput, and long workflow times^15–17^. Here, Ultra-dPCR achieves true single molecule DNA counting with a 6-log dynamic range using a simple and cost-effective workflow without microfluidics. Every sample is partitioned into >30 million droplets, enabling single molecule occupancy via limited dilution, easily extending beyond 1 million DNA molecules (Figure 1).

Ultra-dPCR workflow is simple and utilizes standard molecular biology laboratory tools. Ultra-dPCR mix is centrifuged through a droplet generator directly into PCR strip tubes. Using a unique formulation of emulsion reagents, Ultra-dPCR emulsions are optically transparent, allowing for the use of a 3D imaging technique to rapidly scan each closed PCR tube for DNA-positive droplets. Furthermore, due to single molecule partitioning, Ultra-dPCR enables straightforward multiplexing, avoiding PSC and providing leniency in assay design aspects associated with balancing PCR efficiency across targets. As a proof-of-principle, we demonstrate the use of a 222-plex Ultra-dPCR assay for counting loci of chromosomes 13, 18, and 21 for non-invasive fetal aneuploidy detection, paving way to replace NGS and microarray with a low-cost and rapid workflow in clinical labs.^11,21,22^

## EXPERIMENTAL SECTION

### Ultra-dPCR

PCR reaction mix was prepared using Ultra-dPCR TaqMan Mix (Enumerix) according to the manufacturer’s protocol. The PCR mix was added to the ultra-partitioning droplet generator (Enumerix), outfitted into PCR strip tubes carrying emulsifying reagents (Enumerix) as a strip assembly (Figure 2). The strip assembly was then fitted into an Ultra-dPCR swing bucket (Enumerix) for use in an Eppendorf 5430 centrifuge installed with a swing bucket rotor. For each centrifugation, up to 12 strip assemblies (48 samples) were spun for 20 minutes at 16,000 x g to form Ultra-dPCR emulsions. PCR amplification was performed on Bio-Rad C1000 thermal cyclers. Ultra-dPCR amplified emulsions were scanned via the Ultra-dPCR Imager (Enumerix), followed by 3D image reconstruction to count DNA-positive droplets and measure emulsion volume using a custom software written in C++ and MATLAB. Unless specified, all Ultra-dPCR counts data are normalized to 50 μL input.

**Figure 2.**
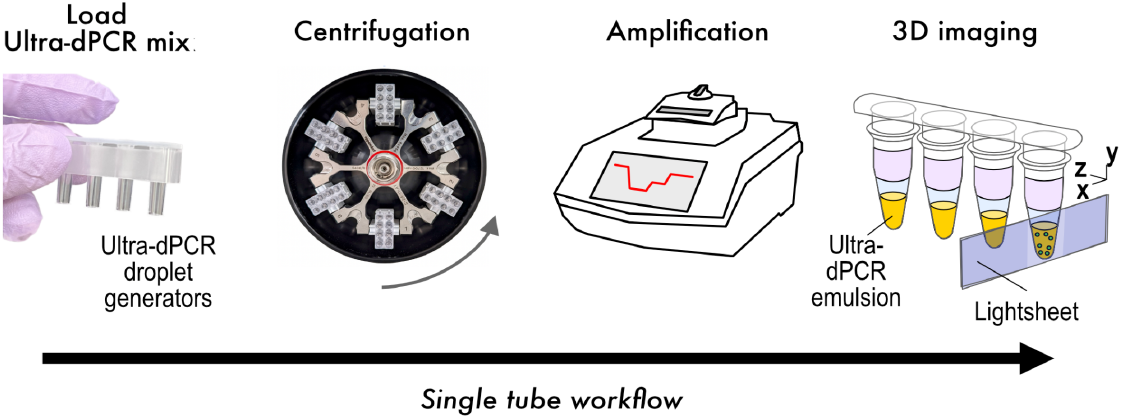
Ultra-dPCR workflow.

### Assay designs

Variations of the CDC SARS-CoV-2 TaqMan assays were used for Ultra-dPCR characterization and benchmarking studies. For 1-color tests, human *RPP30* TaqMan assay was used, and for 2-color tests, human *RPP30* and SARS-CoV-2 *N1* assays were used23. For 4-color tests, human *RPP30*, SARS-CoV-2 *N1*, SARS-CoV-2 *N2*, and an additional *prfA* target from *L. monocytogenes prfA* were used24. The 222-plex NIPT-Ultra-dPCR Taqman assay panel (Enumerix) targeted 74 regions each on chromosomes 13, 18, and 21. The panel was developed based on *in silico* primer design followed by empirical optimization. Primers were designed to target regions outside of common CNVs and SNPs, and with minimal interaction with probes and other primers in the panel. Each individual assay was tested on genomic DNA samples to ensure it produced the correct chromosomal ratios before combining into the final panel. Oligonucleotides, including probes and primers, were purchased from Integrated DNA Technologies.

### Benchmarking against conventional ddPCR

Bio-Rad QX200 2-color ddPCR system was used according to the manufacturer’s instructions at Stanford Genomics. All dPCR reactions used the ddPCR™ Supermix for Probes (Bio-Rad) according to the manufacturer’s instructions. Annealing temperatures for *RPP30* and SARS-CoV-2 *N1* assays were performed using an annealing gradient in a 2-step amplification protocol according to the manufacturer’s instructions. Data analysis was performed using the manufacturer’s Quantasoft software.

### Sample preparation

For SARS-CoV-2 assays, DNA templates were amplified from a cloned plasmid with a full gene insert (Integrated DNA Technologies) using M13 primers, followed by quantification using Ultra-dPCR to obtain DNA concentration. For *prfA*, DNA templates were amplified from synthetic DNA templates using targeted primers, followed by quantification using Ultra-dPCR to obtain DNA concentration. Titration studies were performed using serial dilution with TE Buffer (Teknova). For the fetal aneuploidy assay, contrived DNA samples were generated using nucleosomal DNA from GM12878 (Coriell) and Detroit 532 (ATCC). Nucleosomal DNA was isolated from frozen cell pellets using Zymo Research EZ Nucleosomal DNA Prep Kit and Atlantis dsDNase for cutting and purification. DNA was quantified using Agilent TapeStation D1000 High Sensitivity ScreenTape or cfDNA ScreenTape, followed by quantitation of genome equivalents (GE) by Ultra-dPCR before spiking Detroit 532 (Trisomy 21) DNA into GM12878 to generate contrived DNA mixtures containing various percentages of trisomy 21 DNA. The ploidy of the GM12878 and Detroit 532 were confirmed via NGS as follows: sequencing libraries were generated using NEBNext Ultra II DNA Library Prep Kit for Illumina (New England Biolabs), and libraries were sequenced on a MiSeq (Illumina) with 10% PhiX spike-in. NGS data was analyzed by mapping sequencing reads via CNVkit to determine aneuploidy status11. For cell free DNA (cfDNA), blood was collected in Cell-Free DNA BCT Tubes (Streck) from healthy donors. Cell-free plasma was isolated using the Apostle MiniMax Kit (Beckman Coulter) according to the manufacturer’s protocol. cfDNA was quantified using Agilent TapeStation with the Cell-free DNA ScreenTape Assay.

### Droplet Characterization

Droplet size was measured using brightfield microscopy. Briefly, an Ultra-dPCR emulsion was generated by centrifuging 50 μL PCR emulator mix (absent of DNA template and dNTPs) directly into PCR strip tubes with a spin duration of 20 minutes at 16,000 x g. Droplets were aspirated and diluted to 1:15 with Emulsion Oil (Enumerix). 10 μL of diluted droplets were pipetted into a disposable hemocytometer chip (Incyto DHC-N01) and imaged on a brightfield microscope (AmScope IN200TB). Droplet diameter was quantified with a custom MATLAB script. Packing efficiency of the emulsion was analyzed using 3D lightsheet microscopy with Nile Red (Sigma) stained emulsion oil. Scalability of emulsion generation was evaluated using a 1-color *RPP30* assay where a large Ultra-dPCR master mix was made, followed by emulsion generation with different volume inputs of Ultra-dPCR mixes, and the quantification of total droplets per tube.

## RESULTS AND DISCUSSION

### Single-tube Ultra-dPCR closed workflow

The Ultra-dPCR workflow uses a single closed tube workflow per sample, outlined in Figure 2. Partitioning is achieved via the centrifugation of the PCR mix through disposable droplet generator strip columns and directly into PCR strip tubes (Figure 2). Ultra-dPCR also uses a unique formulation of reagents to form optically clear emulsions with near-100% packing efficiency which contribute to minimal droplet coalescence (Figure 3A and Figure S1A) during physical movement of PCR strip tubes and thermocycling of emulsions. At this packing efficiency, ultra-partitions have no relative motion within the PCR tube and can be directly scanned inside the strip tubes after PCR via 3D lightsheet microscopy in under a 1 minute per channel (Figure 2, Figure 3B). Ultra-dPCR samples can also be stored and rescanned repeatedly (Figure S1B). Throughout this single tube workflow, the Ultra-dPCR mix is contained within a closed system from the point of ultra-partitioning until imaging (Figure 2) with no exchange between samples, thereby significantly reducing the risk of sample-to-sample contamination.

**Figure 3.**
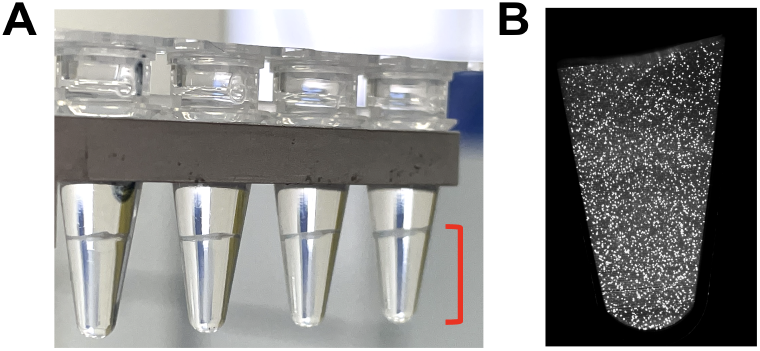
Ultra-partitions characteristics. (A) Ultra-partitions settle in the bottom of the tube (shown in red bracket) and are clear for direct in-tube 3D imaging. (B) A. slice of 3D lightsheet scan of a sample with ∼500,000 counts of human *RPP30* to showcase low DNA occupancy of the system.

### Parallelized ultra-partitioning in a flexible manner

Ultra-dPCR’s ultra-partitions are ∼25-times smaller in volume than another study using centrifugation^24^, and at least 500-times smaller in volume than current commercially available dPCR systems^11,12,21^. The average droplet diameter is 14.1 μm (Figure S2A), equivalent to 1.47 pL in volume, generating >30 million droplets per 50 μL PCR mix from a 20-minute spin using a standard benchtop centrifuge. Ultra-dPCR’s droplet generation system is parallelized and has higher sample throughput, currently with up to 48 samples or >1.5 billion partitions per run.

The Ultra-dPCR droplet generators also accommodate flexible assay volumes, like qPCR. Up to 50 μL of PCR mix can be loaded (**Figure S2B**). For assays with concentrated DNA, a lower reaction volume (i.e., 20 μL) can be used to minimize cost of reagents. For assays with extremely diluted DNA samples (such as cfDNA), as much as 30 μL of template in a 50 μL PCR volume has been successfully processed for use as a downstream DNA quantification tool after cfDNA extraction. This high template-volume feature provides a useful alternative to many other dPCR systems with fixed volumes that are often too low for proper sampling of cfDNA11,25,26.

### Ultra-partitioning with 6-log dynamic range

The partitioning capacity of Ultra-dPCR enables single molecule counting over a broad dynamic range. Even at 1 million DNA copies (or λ = 0.03), Poisson distribution calculates that 98.34% of the partitions have 1 DNA copy (Figure 1). As a proof of principle, we performed a serial dilution experiment of human *RPP30* template targeting a 6-log dynamic range and counted only positive partitions in Ultra-dPCR. We observed linearity in positive partitions throughout the range – where 1 positive partition translates to 1 DNA copy – with low total error throughout this range (Figure 4). To our knowledge, Ultra-dPCR has the highest dynamic range compared to any commercially available dPCR systems while uniquely operating in a single molecule domain, without the need for Poisson correction.

**Figure 4.**
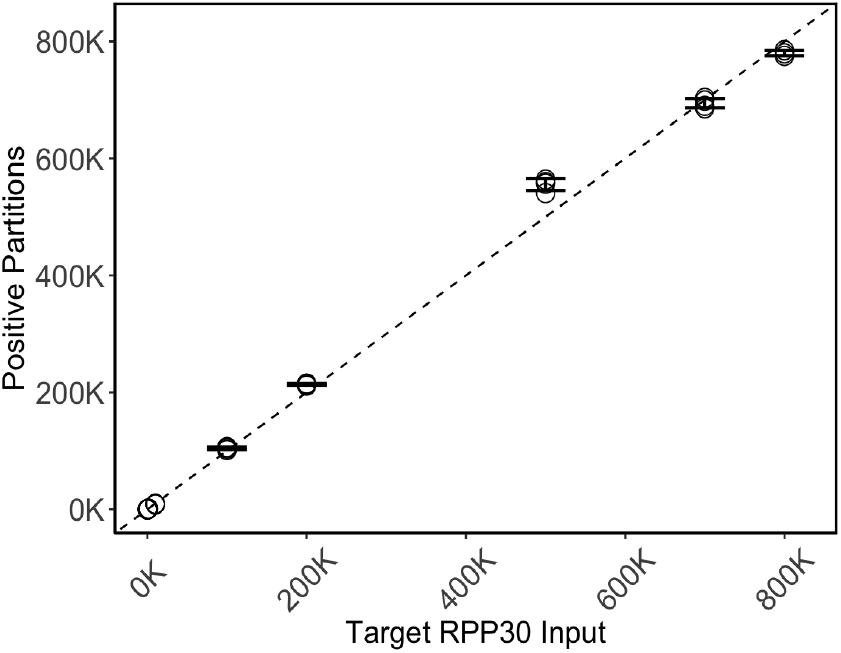
6-log dynamic range on Ultra-dPCR platform. Demonstration of a 6-log dynamic range in serial dilution experiment, measuring DNA-positive partitions vs target *RPP30* input copies. For each dilution, 4 technical replicates are performed, with total error (%CV) represented in the error bars.

### Orthogonal testing for Ultra-dPCR’s linearity and accuracy

Next, we performed orthogonal testing against Bio-Rad QX-200 droplet dPCR (ddPCR). In this study, synthetic SARS-CoV-2 *N1* template was added to the PCR master mix at ∼2K counts per sample prior to dividing the master mix into aliquots to add serially diluted synthetic *RPP30* templates across a 5-log range from 0 to ∼200K copies (denoted S1-S10) (Figure 5, Table S1, Table S2). This multiplex assay was optimized by testing an annealing gradient on both ddPCR and Ultra-dPCR. *N1* amplitude was highly sensitive to the annealing temperature in ddPCR only, with an inverse correlation between annealing temperature and *N1* amplitude (Figure S3A-B), whereas Ultra-dPCR’s *N1* amplitude remained constant (Figure S3C) regardless of annealing temperature. To achieve the best signal-to-noise separation in ddPCR, an annealing temperature of 54°C was used for both systems for direct comparisons of *RPP30* input count accuracy. Using this condition, both ddPCR and Ultra-dPCR had similar *N1* counts throughout the dilution points (Table S1, Table S2). First, we measured the dead volume for each system. In ddPCR, the median number of accepted droplets was 15,620 (Table S1), as compared to the total number of 23,529 droplets generated^12^, equating to ∼33% sample loss. In Ultra-dPCR, via 3D lightsheet imaging, we measured an average flowthrough of ∼53 μL from a target 50 μL PCR mix input. This negligible dead volume suggests that centrifugal forces during partitioning allows near-complete sample utilization. This feature, alongside with up to 30 μL DNA sample input per 50 μL reaction, allows Ultra-dPCR to be particularly useful in applications that require maximal usage of template volume for detecting rare molecules, such as in liquid biopsies.

**Figure 5.**
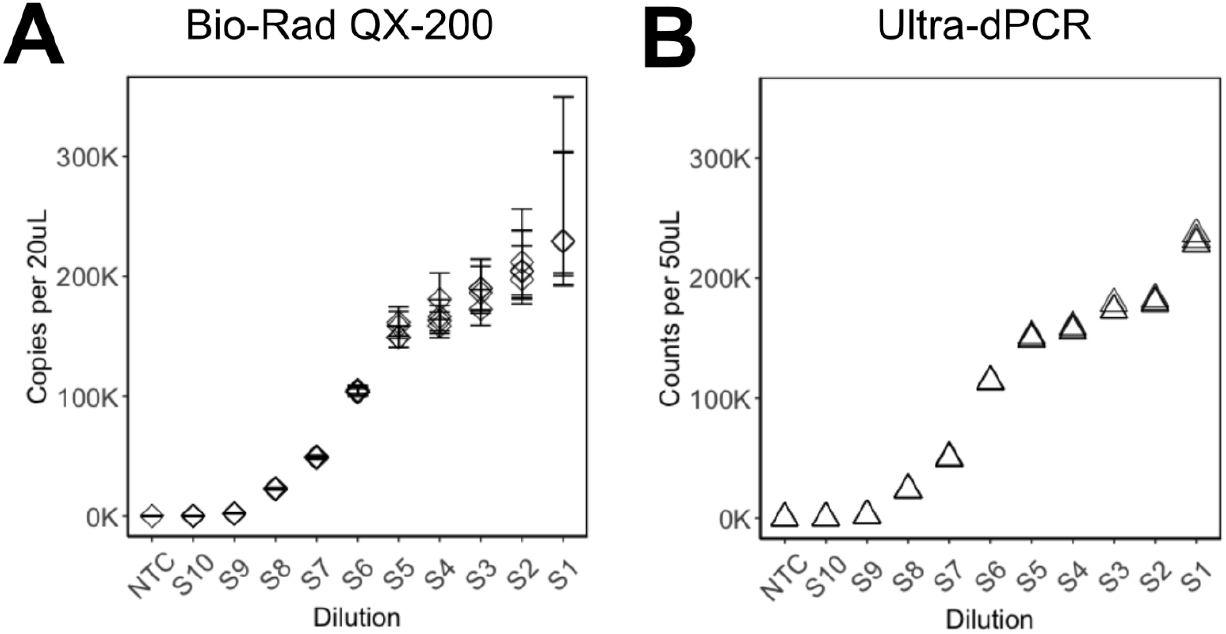
Orthogonal testing between Ultra-dPCR and ddPCR. Scatterplots of *RPP30* counts where the same serially diluted set of samples was tested on (A) ddPCR with Poisson 95% confidence interval (CI) as error bars as indicated by the manufacturer; and (B) Ultra-dPCR with positive partitions normalized to 50uL.

Next, we compared the total *RPP30* copies detected using comparable metrics. For Ultra-dPCR, we reported the number of positive partitions from the entire emulsion, normalized to 50 μL. For ddPCR, we reported the dead-volume adjusted copies per 20 μL from the manufacturer’s software. In low *RPP30* input samples with <50K counts (S7 to S10), we observed similar *RPP30* counts between the two platforms (Figure 5, Table S1, Table S2), suggesting that the two platforms have comparable accuracy at this range.

At higher *RPP30* inputs, such as in beyond 100K counts (S1-S6), counts between ddPCR and Ultra-dPCR began to diverge (Figure 5). Almost all droplets in ddPCR are saturated with DNA, harboring increasing Poisson uncertainty within each technical replicate as *RPP30* input increases (Table S1, Figure 5). The Poisson uncertainty percentages between dilution inputs overlapped, and within the same dilution sample, technical errors (such as from pipetting) were masked by Poisson errors^12^ (Table S1, Figure 5). To derive quantitative accuracy in ddPCR at high range, multiple sample wells are required to distribute DNA counts at a lower occupancy. On the other hand, Ultra-dPCR’s single molecule counting achieves low uncertainty and high resolution in DNA counting. This is exemplified by easily separable *RPP30* counts between tight serial dilutions while also maintaining low and consistent %CVs across this range (Table S2, Figure 5).

The ability to maintain a high dynamic range and high precision counting profile for Ultra-dPCR has distinct advantages. First, for experiments involving DNA quantitation at a high dynamic range, such as gene expression analysis, users do not need to perform further sample dilutions to achieve the desirable precision range, making experimental design more straightforward and with fewer technical errors. Second, in dPCR systems that take advantage of Poisson correction to stretch dynamic range^21^, each positive partition (or negative partition) represents an increasing number of DNA copies within the sample as input increases and thereby reduces the ability to discern DNA count changes. On the other hand, since Ultra-dPCR operates through a single molecule regimen through a 6-log range, every DNA copy is represented proportionally by positive partition counting to achieve high resolution, high precision DNA quantification.

### Ultra-dPCR eliminates partition-specific competition (PSC)

Housing multiple DNA templates within the same partition may be subjected to PSC. This phenomenon occurs when multiple templates are present in the same partition and compete for PCR resources, such as primers, dNTPs, and polymerases, leading to reduction in fluorescence^12,13^. During our orthogonal testing, we observed PSC in ddPCR. As *RPP30* input increased, the amplitude of *N1* decreased (Figure S4A-B). Analysis of the ddPCR 2D plot showed that *RPP30* and *N1* dual-positive droplets had lower *N1* amplitude compared to *N1*-only positive droplets (Figure S4C), likely due to unbalanced PCR efficiencies between the two targets (as demonstrated in the 60°C annealing condition). The decrease in *N1* amplitude led to dropout of DNA counts (Figure S4A), lowering the sensitivity and accuracy of the assay. On the contrary, in Ultra-dPCR, N1 assay amplitude was unaffected with increasing RPP30 inputs (Figure S4D) in both 54°C and 60°C annealing temperature conditions.

The absence of PSC in Ultra-dPCR is attributed to its large partitioning capacity and low DNA occupancy per partition (∼3% with 1 million occupied partitions), enabling virtually every DNA template to be partitioned, amplified, and detected independently from other targets. The elimination of PSCs in Ultra-dPCR allows more straightforward multiplex assay optimization. Multiplex assay design detecting rare molecules in the backdrop of a highly expressed control gene – such as infectious disease and minimal residual disease applications – can be performed more directly without PSC diminishing rare molecule signal. Moreover, single template separation in Ultra-dPCR can aid high order multiplexing assay designs to achieve NGS-like precision DNA counting.

### Ultra-dPCR NIPT assay

In addition to the elimination of PSCs, Ultra-dPCR’s partition capacity also enables new applications beyond standard dPCR, such as NIPT. Fetal aneuploidy screening has become a common form of noninvasive prenatal testing for expectant mothers in the past decade^27^. The most common method of fetal aneuploidy screening relies on counting chromosomal copies using technologies such as NGS or microarrays^28–31^. These tests count copies from different chromosomes in maternal cell-free DNA. Since the fetal fraction in maternal cfDNA is low, technologies that enable high order DNA counting – such as NGS – are used to achieve statistical confidence to detect aneuploidy in as low as 4% fetal fraction conditions^32–34^. At this fetal fraction, these NIPT tests are required to detect chromosomal count differences of 2%. The estimated chromosomal count required to achieve this goal is >220,000 counts for reference chromosomes^35^. The more molecules counted, the higher the statistical confidence for samples with low fetal fractions. With the flexibility of a 6-log counting range coupled with high precision, we hypothesize that Ultra-dPCR can provide a high-performance and cost-effective alternative to NGS for NIPT.

We first determined the average yield of a standard 10 mL blood draw from healthy donors to be ∼10 ng, which is ∼1500 genome equivalents (GEs). To satisfy the 220,000-count requirement for cfDNA samples, at least a 48-plex per chromosome assay is required, which is a ∼5X higher requirement than that for published NIPT dPCR studies^18–20,36^. We tackled this problem via rigorous over-designing of our fetal aneuploidy panel to enable a 74-plex per chromosome, 3-color assay for a total of 222-plex. In total, this is over 10X higher in multiplexing capability than any reported NIPT dPCR assays ^18–20,36^. In this NIPT-Ultra-dPCR assay, each of the chromosomes Chr13, Chr18, and Chr21 are targeted with a set of optimized chromosome-specific probes having different colored fluorophores, which are compatible with the Ultra-dPCR emulsion system and 3D imager, and which achieve high signal amplitude and low background across various DNA input. We validated each of the 222-plex individually to ensure that they produced the expected counts to the expected chromosome with no cross-reaction to another chromosome. The total counts from the 74-plex assay also correlated closely to the healthy donor cfDNA input, indicating that the assay is highly efficient (Figure S5A). We have also performed thermal cycler variability testing to ensure reproducible results in chromosomal ratios across 3 available thermal cyclers (Figure S5B).

We generated contrived DNA samples mimicking different percentages of trisomy 21 (%T21) at 0%, 4%, 6%, and 10% in a background of euploid DNA. We tested them at conservative target of ∼1667 GE or 11ng input. Summary of this experiment is provided in Table 1. When considering Ultra-dPCR absolute molecule counts, we observed 2.591 to 3.811% CV between technical replicates, which was likely due to technical error between pipetting steps. However, when analyzing chromosome ratios Chr21/13, Chr21/18, and Chr13/18, we observed a much lower CV of 0.319 to 0.647% between replicates. Utilizing this 3-color assay, we used both Chr13 and Chr18 as reference chromosomes to detect minute Chr21 elevation in the contrived samples. When analyzing Chr21/Chr13 and Chr21/Chr18 ratios, we observed a distinct shift in ratios as %T21 increased, (Figure 5A-B), while Chr13/18 ratio remained similar as expected (Figure 5C).

**Table 1.** NIPT-Ultra-dPCR result summary.

| %Trisomy 21 | Metric | Chr13<br>Counts | Chr21<br>Counts | Chr18<br>Counts | Chr21/13<br>Ratio | Chr21/18<br>Ratio | Chr13/18<br>Ratio |
| --- | --- | --- | --- | --- | --- | --- | --- |
| 0 | Mean | 272589 | 284675 | 270602 | 1.044 | 1.052 | 1.007 |
|  | (%CV) | (2.798%) | (2.644%) | (2.696%) | (0.343%) | (0.377%) | (0.319%) |
| 4 | Mean | 276641 | 293851 | 273524 | 1.062 | 1.074 | 1.011 |
|  | (%CV) | (3.070%) | (3.210%) | (3.089%) | (0.393%) | (0.378%) | (0.397%) |
| 6 | Mean | 262783 | 281465 | 259506 | 1.071 | 1.085 | 1.013 |
|  | (%CV) | (3.630%) | (3.811%) | (3.777%) | (0.402%) | (0.532%) | (0.647%) |
| 10 | Mean | 266260 | 289984 | 263157 | 1.089 | 1.102 | 1.012 |
|  | (%CV) | (2.591%) | (2.808%) | (2.692%) | (0.456%) | (0.403%) | (0.359%) |

Next, we performed receiver operating characteristic (ROC) analysis to identify optimal threshold to different T21-spiked versus 0% T21 contrived samples. Unlike previous NIPT dPCR attempts with only one reference chromosome via a 2-color test^6,19,20^, we evaluated trisomy calls using two reference chromosomes. Through this analysis, both Chr21/13 and Chr21/18 ratio analysis yielded 100% sensitivity, specificity, and accuracy across all tested %T21 conditions (cutoff values are 1.054 for Chr21/Chr13 and 1.066 for Chr21/Chr18, Figure 5C-D). To our knowledge, this is the first published dataset to distinguish a DNA mixture containing 4% T21 using dPCR without sample enrichment technologies.

dPCR has long been touted as a method of determining fetal aneuploidy but has not yet been widely adopted clinically due to technical limitations^8,19,20,36,37^. First, no commercially available dPCR system can meet the precision requirement due to low partitioning capacity. Multiple groups have tried splitting a sample into multiple sample wells to add more partitions^19^, or perform a series of fetal cfDNA pre-enrichment of fetal cfDNA^36^, adding cost and complexity to the workflow. Splitting a sample across multiple wells or pre-enrichment of fetal cfDNA can also introduce additional errors that may skew true chromosomal ratios. More recently, a group has added more virtual partitions within the color space using a limited primer approach combined with complex analysis techniques to discern >5% T21 samples ^18^. In contrast, even in its proof-of-concept phase, Ultra-dPCR already provides the quantitative precision to differentiate chromosomal count differences as low as 2%. This precision extends beyond NIPT and can be applied to discern minute gene expression changes that qPCR and dPCR cannot currently achieve.

Second, no non-NGS or non-microarray assay sufficiently high multiplexing capabilities to interrogate enough genomic loci to achieve the counting requirement using limited cell-free DNA available from a typical blood draw volume (∼10 mL whole blood). Unlike NGS where bioinformatics pipelines can remove non-specific sequencing reads, dPCR assays must be developed at a high rigor to have low false-positive and false-negative signals for proper droplet counting. Part of this achievement with our 222-plex assay is due to Ultra-dPCR’s single molecule amplification regimen with no interference from other templates. In contrast, with low-partitioning systems, as DNA occupancy (or λ) increases, PSC becomes a larger problem, requiring arduous assay development work to obtain enough DNA counts to achieve the precision required for NIPT. This may explain why all published NIPT-dPCR assays utilize designs with low multiplexing capability^18–20,36^, requiring a much higher DNA input, or pre-enrich amplicons^18^ than what is commonly extracted from peripheral blood. Altogether, Ultra-dPCR’s unique ultra-partitioning method enables unprecedented multiplexing capability with high precision.

## CONCLUSIONS

Ultra-dPCR is a novel platform that re-imagines DNA counting. With >30 million partitions, DNA molecules are partitioned into a single molecule domain through a 6-log dynamic range (Figure 1), providing high resolution DNA counting beyond current dPCR capabilities. Every positive partition translates to one DNA copy, enabling straightforward data interpretation and high precision data without error correction. In addition, when single DNA templates are isolated for amplification, there is no PSC, easing the need to perform rigorous assay design to balance PCR efficiency for multiplexing.

In addition to standard dPCR applications, Ultra-dPCR can be extended to research areas where a high dynamic range and/or high precision counting is required. One such example is in RNA expression where the dynamic range is orders of magnitudes larger than DNA and can vary rapidly through transcriptional and posttranscriptional regulation. In these applications, dPCR is not commonly used, instead users opt for a workflow where RNA markers are identified using NGS, and experimentally verified and validated using qPCR. While qPCR is simple with high dynamic range, it comes with its limitations. qPCR requires a standard curve or housekeeping gene for DNA quantification, and because signal is quantified after every PCR cycle, the precision is only 2-fold. Increasing qPCR resolution is possible and is supported by a study where as many as 18 qPCR replicates are required to discern a 25% difference in DNA counts^38^. However, this type of qPCR experimental design is costly, laborious, consumes a lot of the DNA sample, and generally not practical. On the other hand, Ultra-dPCR offers a similar dynamic range in a simple workflow and can discern a DNA count difference as low as 2% as demonstrated in our NIPT analytical study (Figure 6). With such high resolution and a precision profile that rivals NGS but at a fraction of the cost, Ultra-dPCR can be positioned as a powerful and practical validation partner tool for biomarker research.

**Figure 6.**
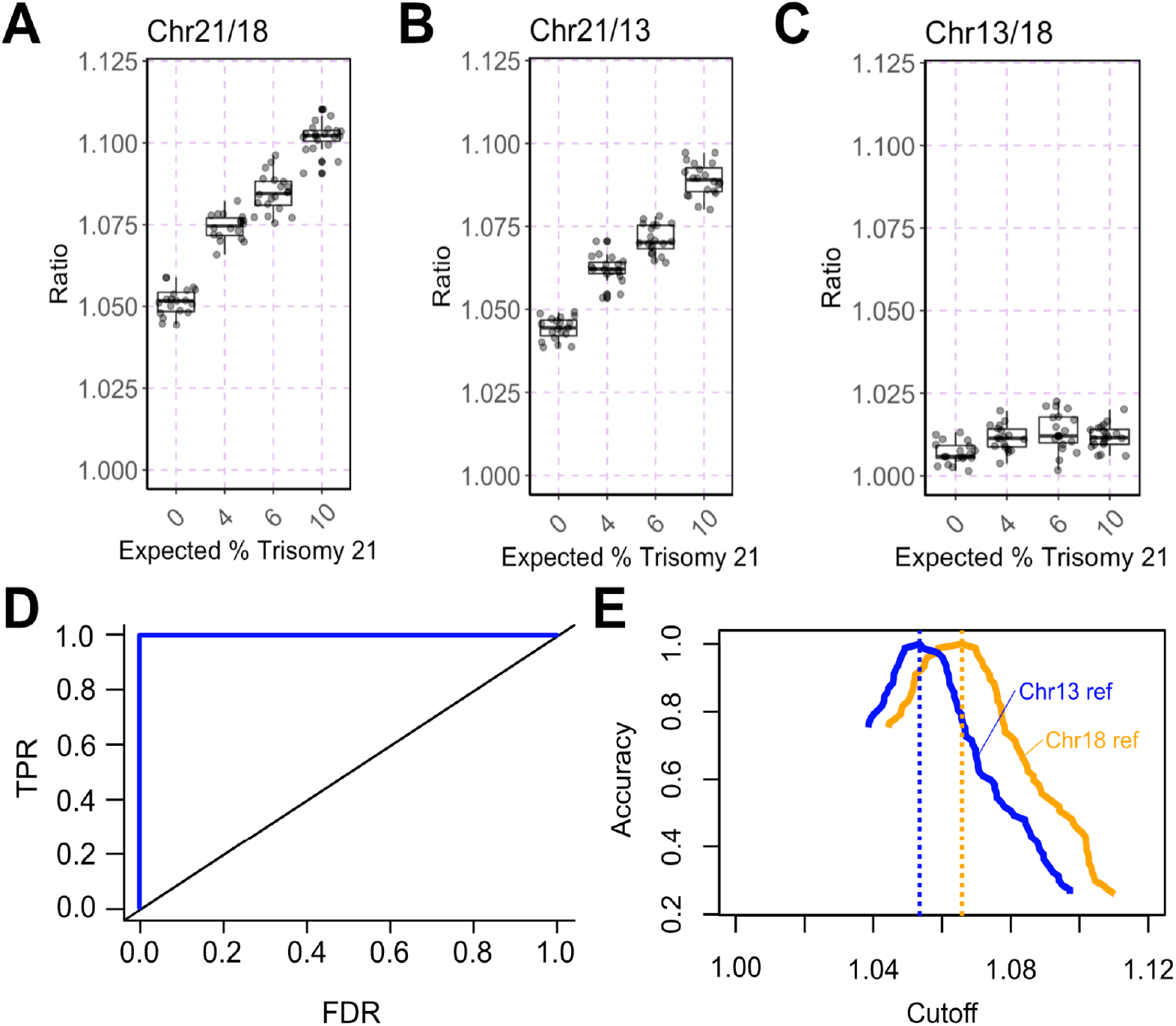
Ultra-dPCR NIPT assay analytical study. (A-C) Boxplot of different chromosomal ratios. Chr21/18 and Chr21/13 are expected to increase with more %T21 spiked in, and Chr13/18 is expected to remain constant. (D) ROC curve using all spiked %T21 samples compared to 0% to determine optimal threshold to separate control versus T21-spiked samples. (E) Accuracy plot vs cutoff where blue line represents data using Chr13 as reference chromosome, and orange line represents data using Chr18 as reference chromosome. Dotted lines represent their respective optimal cutoff to maximize accuracy of the assay.

Beyond the research setting, Ultra-dPCR provides a swift workflow in high precision clinical assays. As a proof-of-concept, we developed an industry-leading 222-plex NIPT assay for fetal aneuploidy. This NIPT assay can likely be further optimized to include additional chromosomes and genes to detect sex aneuploidies and single-gene disorders. While this assay is still at its early stages, the process of Ultra-dPCR is simple and can enable a potential single-day workflow at dPCR-like cost without the need for heavy computational biology support, paving the way for NIPT to be decentralized to hospital labs and beyond first-world countries.

## ASSOCIATED CONTENT

### Supporting Information

The Supporting Information is available free of charge on the ACS Publications website.

Supplementary figures and table provided in a PDF format.

## AUTHOR INFORMATION

### Author Contributions

‡These authors contributed equally. E.Y.S., J.H.L., S.L., S.P.A.F., and H.C.F. conceptualized, designed, and developed the Ultra-dPCR technology and research. H.G.L., J.S., V.K.R., and C.K.L. performed the research. E.Y.S., J.H.L., S.L., H.G.L., J.S. analyzed the data. E.Y.S wrote the manuscript.

### Notes

Competing Interest Statement: The authors declare the following competing financial interest(s): E.Y.S., J.H.L., S.L., H.G.L., J.S., V.K.R., C.K.L., S.P.A.F., H.C.F. are employees of Enumerix, Inc., a company that commercializes DNA counting technologies. Enumerix, Inc., solely funded this study. In addition, E.Y.S., J.H.L., S.L., S.P.A.F., H.C.F. are inventors on patents and patent applications relating to methods described in this manuscript.

## ACKNOWLEDGMENT

We thank Mr. Ari Chaney, Ms. Kat George, Mr. Ivan Wong, Ms. Camille Barker, and Mr. Billy Chan for their support of this research.

## SUPPORTING INFORMATION TO

**Figure S1.**
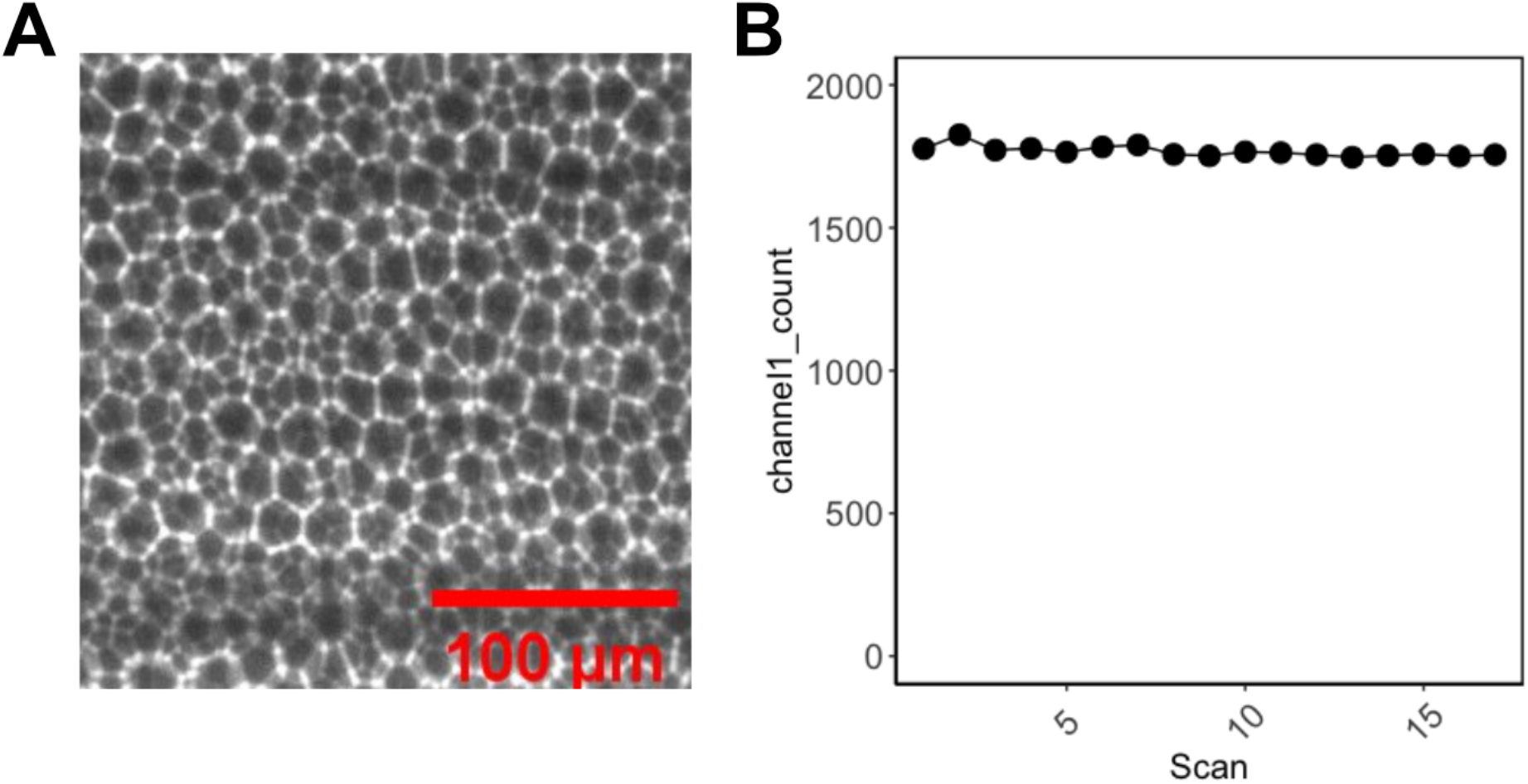
Ultra-partitions have high packing efficiency and are immobilized. (A) Image from lightsheet microscopy of ultra-partitions where Nile Red oil stain is spiked into the emulsion oil to visualize droplet shells. The centrifugation step during ultra-partitioning enables droplets to be densely packed, forming a viscous gel that immobilizes the emulsion to be suitable for 3D imaging and subsequent storage. (B) Ultra-dPCR emulsion stability test. A sample with amplified *prfA* template was repeatedly imaged 17 times over the course of 7 months, showing low variability in DNA counts.

**Figure S2.**
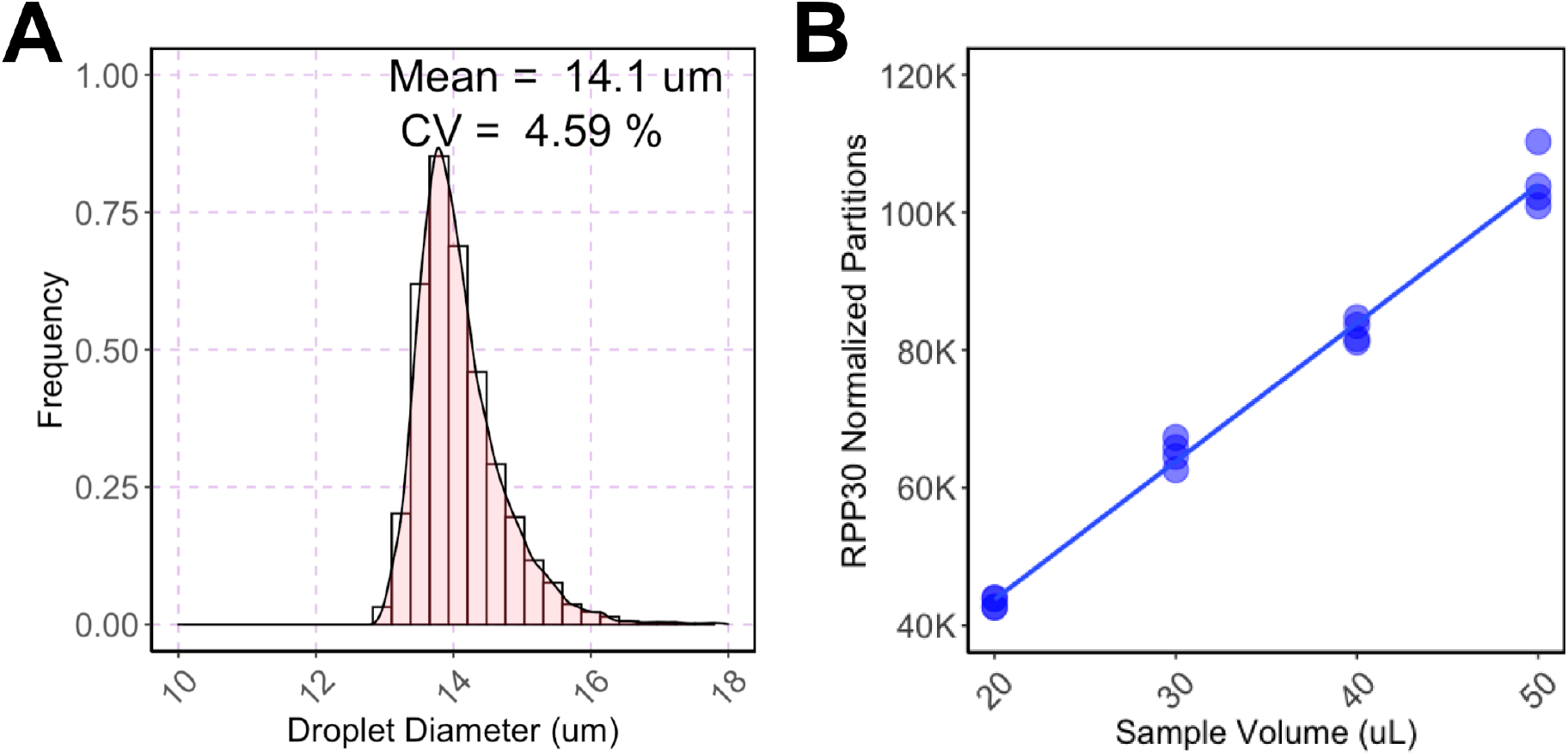
Ultra-dPCR emulsion characterization. (A) Droplet diameter of ultra-partitions, analyzed via brightfield microscopy and MATLAB. (B) Volume titration using the same PCR mix to amplify human *RPP30* amplicon. Each volume has 4 technical replicates with a linear regression fitted (Adjusted R2 = 0.9621).

**Figure S3.**
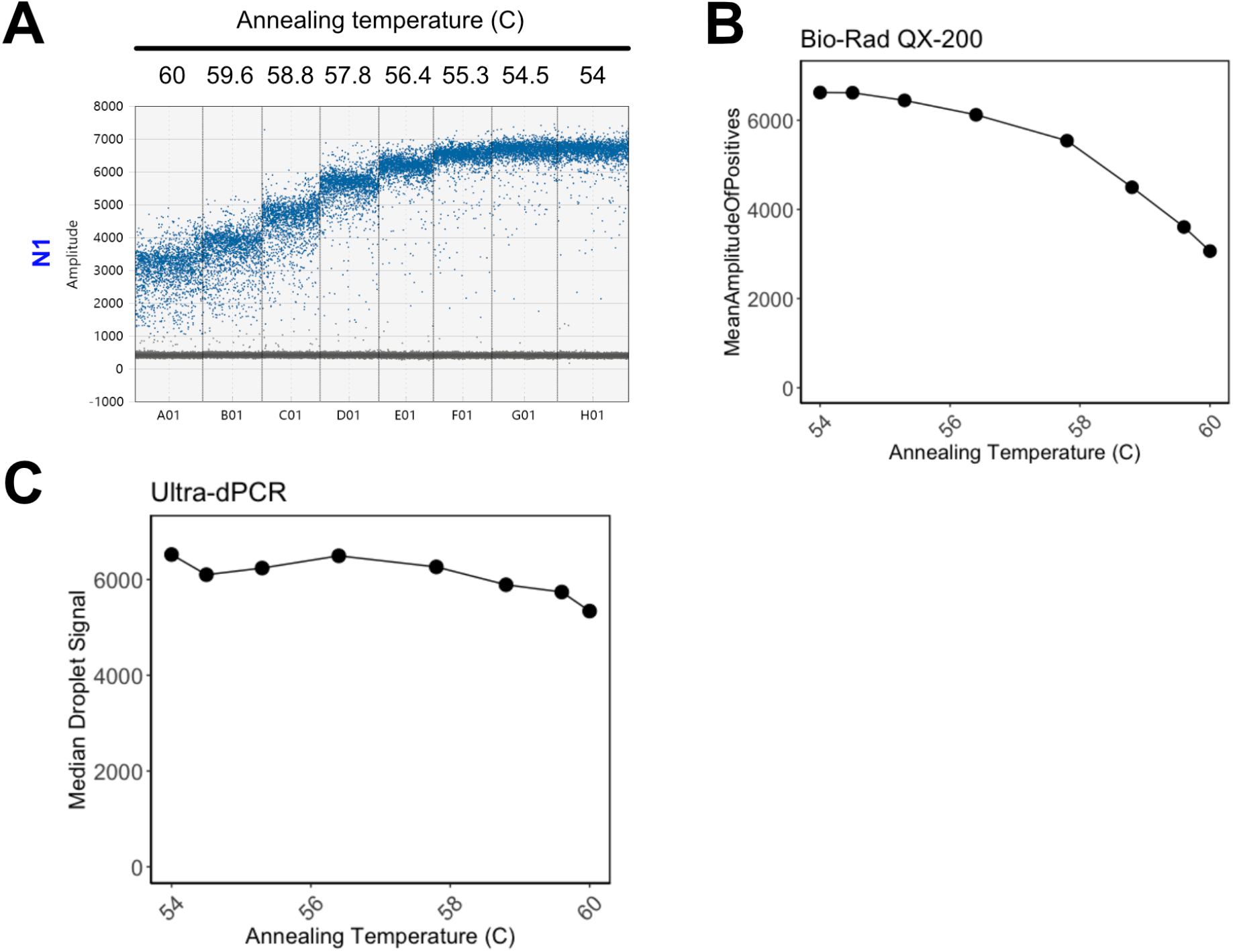
Assay optimization for orthogonal testing. (A) 1D amplitude plot representing data from annealing temperature gradient on QX-200 of SARS-CoV-2 N1 amplicon. (B) Mean amplitude of positive droplets versus annealing temperature as provided by the QuantaSoft software. (C) Median amplitude of *RPP30*-positive droplets measured by Ultra-dPCR.

**Figure S4.**
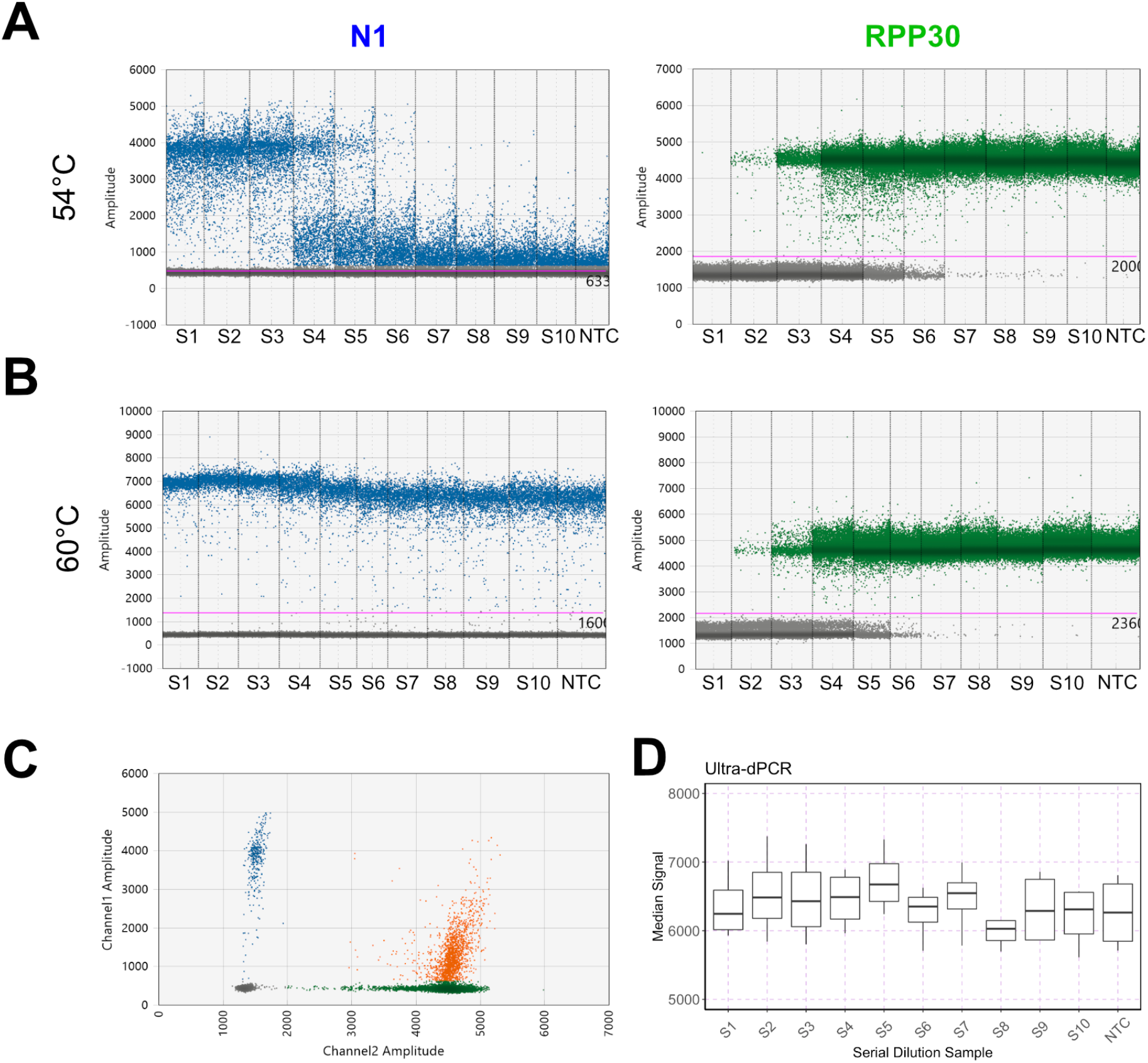
Ultra-dPCR is immune to partition-specific competition. (A) 1D amplitude of SARS-CoV-2 *N1* (left, channel1) and *RPP30* (right, channel2) from QX-200 with different *RPP30* dilutions at 54°C annealing temperature. (B) 1D amplitude of SARS-CoV-2 *N1* (left, channel1) and *RPP30* (right, channel2) from QX-200 with different *RPP30* dilutions at 60°C annealing temperature. (C) 2D amplitude of a sample in (A) where orange points denote droplets that are channel1+ and channel2+, blue points denote channel1+ only, green points denote channel2+ only, and grey points denote double negative droplets. (D) Median signal of positive droplets of SARS-CoV-2 N1 in Ultra-dPCR displayed as a boxplot for 4 technical replicates per serial dilution.

**Figure S5.**
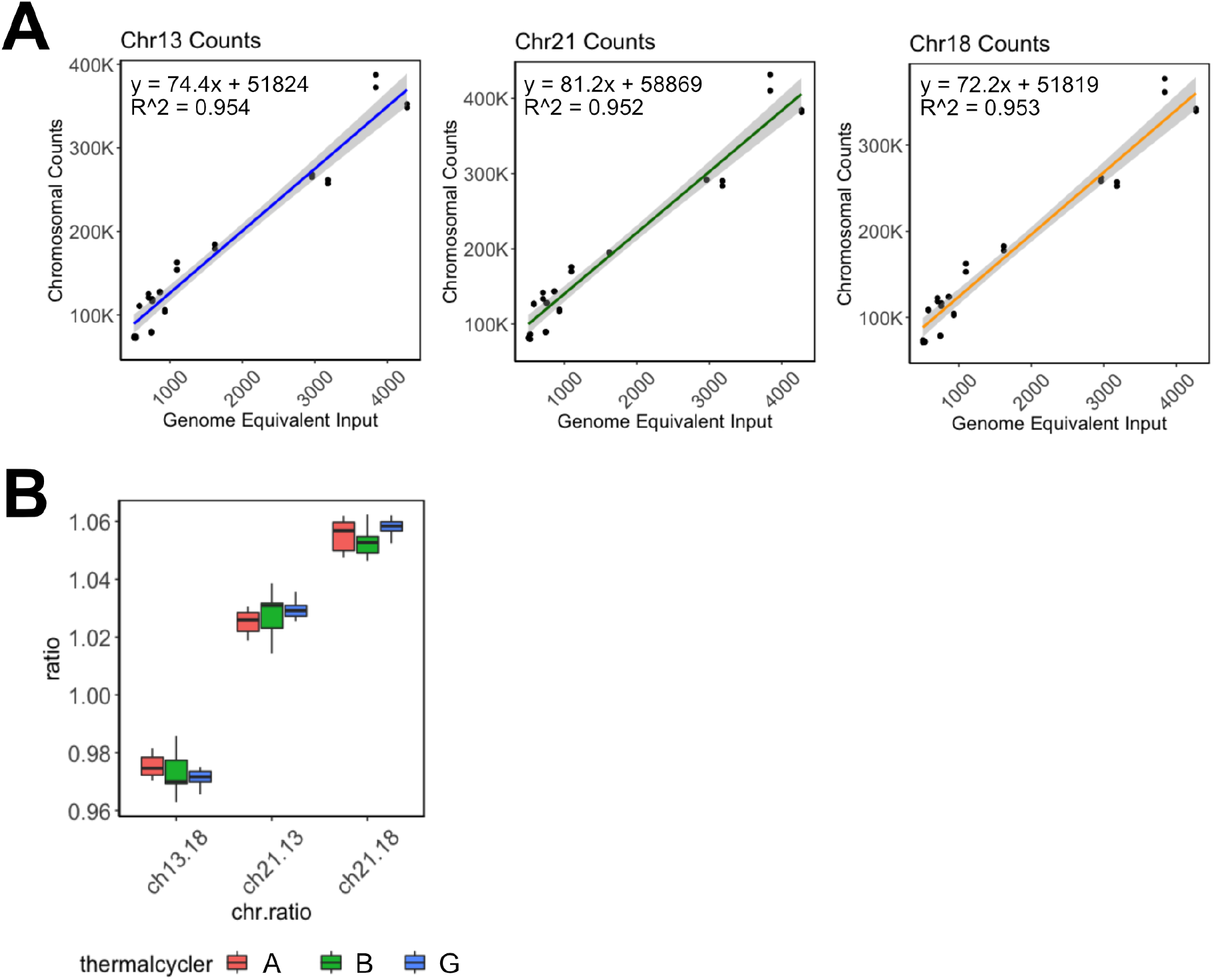
Ultra-dPCR NIPT assay characterization. (A) cfDNA samples from healthy donors measured using Ultra-dPCR NIPT assay versus TapeStation cfDNA ScreenTape (ng of cfDNA converted to genome equivalent (GE), based on a 6.6 pg/GE conversion factor). Two 10 mL blood tubes were obtained per donor and for each blood tube, extracted cfDNA was divided into 2 Ultra-dPCR reactions. For the Ultra-dPCR NIPT assay, Chr13, Chr18, and Chr21 counts are measured and compared independently to TapeStation data, fitted with a linear regression. Slope of chromosomal counts to GE is between 72.2-81.2, which is close to the expected increase of 74-plex per chromosome of the assay, suggesting high efficiency in assay performance. (B) Ultra-dPCR NIPT assay is optimized to have low variability between thermal cyclers, as shown in this thermal cycler reproducibility study measuring the differences in chromosomal ratios.

**Table S1.** Orthogonal testing data on Bio-Rad QX-200.

| Well | Dilution | Target | Conc(copies/μL) | Copies/20μLWell | PoissonConfMax | PoissonConfMin | Accepted Droplets | Positives | Negatives | Ch1+Ch2+ | Ch1+Ch2- | Ch1-Ch2+ |
| --- | --- | --- | --- | --- | --- | --- | --- | --- | --- | --- | --- | --- |
| A01 | NTC | N1-FAM | 127 | 2545 | 134 | 121 | 15276 | 1566 | 13710 | 0 | 1566 | 0 |
| B01 | NTC | N1-FAM | 124 | 2489 | 131 | 118 | 12930 | 1298 | 11632 | 0 | 1298 | 0 |
| C01 | NTC | N1-FAM | 119 | 2377 | 125 | 113 | 16944 | 1628 | 15316 | 0 | 1628 | 0 |
| D01 | NTC | N1-FAM | 121 | 2430 | 127 | 116 | 16809 | 1649 | 15160 | 0 | 1649 | 2 |
| A01 | NTC | RPP30-HEX | 0 | 0 | 0.231 | 0 | 15276 | 0 | 15276 | 0 | 1566 | 0 |
| B01 | NTC | RPP30-HEX | 0 | 0 | 0.273 | 0 | 12930 | 0 | 12930 | 0 | 1298 | 0 |
| C01 | NTC | RPP30-HEX | 0 | 0 | 0.208 | 0 | 16944 | 0 | 16944 | 0 | 1628 | 0 |
| D01 | NTC | RPP30-HEX | 0.14 | 2.8 | 0.448 | 0.0212 | 16809 | 2 | 16807 | 0 | 1649 | 2 |
| A02 | S10 | N1-FAM | 128 | 2553 | 134 | 121 | 14100 | 1450 | 12650 | 6 | 1444 | 87 |
| B02 | S10 | N1-FAM | 122 | 2448 | 129 | 116 | 14402 | 1423 | 12979 | 4 | 1419 | 88 |
| C02 | S10 | N1-FAM | 116 | 2311 | 121 | 110 | 15619 | 1461 | 14158 | 12 | 1449 | 101 |
| D02 | S10 | N1-FAM | 123 | 2467 | 130 | 117 | 15620 | 1555 | 14065 | 17 | 1538 | 88 |
| A02 | S10 | RPP30-HEX | 7.79 | 156 | 9.49 | 6.3 | 14100 | 93 | 14007 | 6 | 1444 | 87 |
| B02 | S10 | RPP30-HEX | 7.54 | 151 | 9.19 | 6.1 | 14402 | 92 | 14310 | 4 | 1419 | 88 |
| C02 | S10 | RPP30-HEX | 8.54 | 171 | 10.1 | 6.97 | 15619 | 113 | 15506 | 12 | 1449 | 101 |
| D02 | S10 | RPP30-HEX | 7.94 | 159 | 9.45 | 6.42 | 15620 | 105 | 15515 | 17 | 1538 | 88 |
| A03 | S9 | N1-FAM | 122 | 2444 | 129 | 116 | 14598 | 1440 | 13158 | 122 | 1318 | 1104 |
| B03 | S9 | N1-FAM | 123 | 2458 | 129 | 117 | 14790 | 1467 | 13323 | 126 | 1341 | 1104 |
| C03 | S9 | N1-FAM | 122 | 2442 | 129 | 116 | 14020 | 1382 | 12638 | 104 | 1278 | 983 |
| D03 | S9 | N1-FAM | 123 | 2450 | 129 | 116 | 15865 | 1569 | 14296 | 153 | 1416 | 1171 |
| A03 | S9 | RPP30-HEX | 103 | 2064 | 109 | 97.4 | 14598 | 1226 | 13372 | 122 | 1318 | 1104 |
| B03 | S9 | RPP30-HEX | 102 | 2043 | 108 | 96.5 | 14790 | 1230 | 13560 | 126 | 1341 | 1104 |
| C03 | S9 | RPP30-HEX | 94.9 | 1899 | 101 | 89.3 | 14020 | 1087 | 12933 | 104 | 1278 | 983 |
| D03 | S9 | RPP30-HEX | 103 | 2050 | 108 | 97 | 15865 | 1324 | 14541 | 153 | 1416 | 1171 |
| A04 | S8 | N1-FAM | 133 | 2655 | 139 | 126 | 15569 | 1661 | 13908 | 1053 | 608 | 8759 |
| B04 | S8 | N1-FAM | 124 | 2474 | 130 | 117 | 14746 | 1472 | 13274 | 899 | 573 | 8116 |
| C04 | S8 | N1-FAM | 123 | 2470 | 130 | 117 | 15888 | 1583 | 14305 | 1004 | 579 | 8921 |
| D04 | S8 | N1-FAM | 122 | 2445 | 128 | 116 | 15815 | 1561 | 14254 | 938 | 623 | 8708 |
| A04 | S8 | RPP30-HEX | 1170 | 23409 | 1195 | 1147 | 15569 | 9812 | 5757 | 1053 | 608 | 8759 |
| B04 | S8 | RPP30-HEX | 1112 | 22237 | 1136 | 1088 | 14746 | 9015 | 5731 | 899 | 573 | 8116 |
| C04 | S8 | RPP30-HEX | 1153 | 23059 | 1177 | 1130 | 15888 | 9925 | 5963 | 1004 | 579 | 8921 |
| D04 | S8 | RPP30-HEX | 1108 | 22151 | 1131 | 1085 | 15815 | 9646 | 6169 | 938 | 623 | 8708 |
| A05 | S7 | N1-FAM | 121 | 2415 | 127 | 115 | 16628 | 1622 | 15006 | 1430 | 192 | 13201 |
| B05 | S7 | N1-FAM | 116 | 2330 | 123 | 110 | 13281 | 1252 | 12029 | 1090 | 162 | 10543 |
| C05 | S7 | N1-FAM | 118 | 2367 | 124 | 112 | 15841 | 1516 | 14325 | 1315 | 201 | 12472 |
| D05 | S7 | N1-FAM | 116 | 2321 | 122 | 110 | 16647 | 1564 | 15083 | 1369 | 195 | 13252 |
| A05 | S7 | RPP30-HEX | 2493 | 49869 | 2543 | 2446 | 16628 | 14631 | 1997 | 1430 | 192 | 13201 |
| B05 | S7 | RPP30-HEX | 2455 | 49101 | 2509 | 2403 | 13281 | 11633 | 1648 | 1090 | 162 | 10543 |
| C05 | S7 | RPP30-HEX | 2403 | 48066 | 2452 | 2357 | 15841 | 13787 | 2054 | 1315 | 201 | 12472 |
| D05 | S7 | RPP30-HEX | 2478 | 49557 | 2527 | 2431 | 16647 | 14621 | 2026 | 1369 | 195 | 13252 |
| A06 | S6 | N1-FAM | 125 | 2499 | 131 | 119 | 16325 | 1645 | 14680 | 1627 | 18 | 14494 |
| B06 | S6 | N1-FAM | 116 | 2317 | 123 | 109 | 11133 | 1044 | 10089 | 1029 | 15 | 9970 |
| C06 | S6 | N1-FAM | 116 | 2313 | 122 | 110 | 15008 | 1405 | 13603 | 1384 | 21 | 13443 |
| D06 | S6 | N1-FAM | 118 | 2354 | 124 | 112 | 15461 | 1472 | 13989 | 1465 | 7 | 13820 |
| A06 | S6 | RPP30-HEX | 5156 | 103114 | 5328 | 5005 | 16325 | 16121 | 204 | 1627 | 18 | 14494 |
| B06 | S6 | RPP30-HEX | 5200 | 103996 | 5417 | 5017 | 11133 | 10999 | 134 | 1029 | 15 | 9970 |
| C06 | S6 | RPP30-HEX | 5197 | 103949 | 5381 | 5038 | 15008 | 14827 | 181 | 1384 | 21 | 13443 |
| D06 | S6 | RPP30-HEX | 5265 | 105308 | 5452 | 5104 | 15461 | 15285 | 176 | 1465 | 7 | 13820 |
| A07 | S5 | N1-FAM | 121 | 2422 | 127 | 115 | 15241 | 1491 | 13750 | 1488 | 3 | 13726 |
| B07 | S5 | N1-FAM | 120 | 2409 | 127 | 114 | 14357 | 1397 | 12960 | 1396 | 1 | 12946 |
| C07 | S5 | N1-FAM | 122 | 2432 | 128 | 115 | 15315 | 1504 | 13811 | 1498 | 6 | 13790 |
| D07 | S5 | N1-FAM | 118 | 2362 | 124 | 112 | 14387 | 1374 | 13013 | 1372 | 2 | 12998 |
| A07 | S5 | RPP30-HEX | 7454 | 149080 | 7929 | 7035 | 15241 | 15214 | 27 | 1488 | 3 | 13726 |
| B07 | S5 | RPP30-HEX | 8075 | 161505 | 8729 | 7523 | 14357 | 14342 | 15 | 1396 | 1 | 12946 |
| C07 | S5 | RPP30-HEX | 7460 | 149194 | 7935 | 7041 | 15315 | 15288 | 27 | 1498 | 6 | 13790 |
| D07 | S5 | RPP30-HEX | 7930 | 158609 | 8541 | 7410 | 14387 | 14370 | 17 | 1372 | 2 | 12998 |
| A08 | S4 | N1-FAM | 117 | 2337 | 123 | 111 | 16213 | 1533 | 14680 | 1530 | 3 | 14664 |
| B08 | S4 | N1-FAM | 121 | 2421 | 128 | 114 | 13069 | 1278 | 11791 | 1277 | 1 | 11786 |
| C08 | S4 | N1-FAM | 121 | 2413 | 127 | 115 | 16435 | 1602 | 14833 | 1602 | 0 | 14817 |
| D08 | S4 | N1-FAM | 129 | 2579 | 135 | 123 | 15288 | 1587 | 13701 | 1584 | 3 | 13691 |
| A08 | S4 | RPP30-HEX | 7940 | 158803 | 8515 | 7446 | 16213 | 16194 | 19 | 1530 | 3 | 14664 |
| B08 | S4 | RPP30-HEX | 9043 | 180853 | 10145 | 8202 | 13069 | 13063 | 6 | 1277 | 1 | 11786 |
| C08 | S4 | RPP30-HEX | 8158 | 163167 | 8790 | 7623 | 16435 | 16419 | 16 | 1602 | 0 | 14817 |
| D08 | S4 | RPP30-HEX | 8317 | 166350 | 9026 | 7728 | 15288 | 15275 | 13 | 1584 | 3 | 13691 |
| A09 | S3 | N1-FAM | 112 | 2233 | 117 | 106 | 16048 | 1453 | 14595 | 1452 | 1 | 14591 |
| B09 | S3 | N1-FAM | 117 | 2331 | 122 | 111 | 16473 | 1554 | 14919 | 1551 | 3 | 14916 |
| C09 | S3 | N1-FAM | 125 | 2496 | 131 | 119 | 15319 | 1542 | 13777 | 1540 | 2 | 13769 |
| D09 | S3 | N1-FAM | 129 | 2571 | 135 | 122 | 16078 | 1664 | 14414 | 1664 | 0 | 14409 |
| A09 | S3 | RPP30-HEX | 9499 | 189974 | 10728 | 8586 | 16048 | 16043 | 5 | 1452 | 1 | 14591 |
| B09 | S3 | RPP30-HEX | 9315 | 186299 | 10418 | 8474 | 16473 | 16467 | 6 | 1551 | 3 | 14916 |
| C09 | S3 | RPP30-HEX | 8629 | 172571 | 9449 | 7963 | 15319 | 15309 | 10 | 1540 | 2 | 13769 |
| D09 | S3 | RPP30-HEX | 9501 | 190018 | 10731 | 8588 | 16078 | 16073 | 5 | 1664 | 0 | 14409 |
| A10 | S2 | N1-FAM | 126 | 2517 | 132 | 120 | 16315 | 1655 | 14660 | 1655 | 0 | 14658 |
| B10 | S2 | N1-FAM | 124 | 2476 | 130 | 118 | 17460 | 1744 | 15716 | 1743 | 1 | 15714 |
| C10 | S2 | N1-FAM | 125 | 2493 | 130 | 119 | 17594 | 1769 | 15825 | 1769 | 0 | 15821 |
| D10 | S2 | N1-FAM | 122 | 2442 | 128 | 116 | 18256 | 1800 | 16456 | 1800 | 0 | 16453 |
| A10 | S2 | RPP30-HEX | 10596 | 211922 | 12816 | 9227 | 16315 | 16313 | 2 | 1655 | 0 | 14658 |
| B10 | S2 | RPP30-HEX | 10199 | 203978 | 11891 | 9052 | 17460 | 17457 | 3 | 1743 | 1 | 15714 |
| C10 | S2 | RPP30-HEX | 9869 | 197389 | 11279 | 8860 | 17594 | 17590 | 4 | 1769 | 0 | 15821 |
| D10 | S2 | RPP30-HEX | 10251 | 205027 | 11943 | 9105 | 18256 | 18253 | 3 | 1800 | 0 | 16453 |
| A11 | S1 | N1-FAM | 124 | 2477 | 130 | 118 | 15101 | 1509 | 13592 | 1509 | 0 | 13592 |
| B11 | S1 | N1-FAM | 120 | 2396 | 126 | 114 | 17505 | 1695 | 15810 | 1695 | 0 | 15809 |
| C11 | S1 | N1-FAM | 120 | 2402 | 126 | 114 | 16733 | 1624 | 15109 | 1624 | 0 | 15109 |
| D11 | S1 | N1-FAM | 123 | 2461 | 129 | 117 | 16773 | 1666 | 15107 | 1666 | 0 | 15106 |
| A11 | S1 | RPP30-HEX | 1000000 | 20000000 | 1000000 | 10030 | 15101 | 15101 | 0 | 1509 | 0 | 13592 |
| B11 | S1 | RPP30-HEX | 11494 | 229888 | 15224 | 9655 | 17505 | 17504 | 1 | 1695 | 0 | 15809 |
| C11 | S1 | RPP30-HEX | 1000000 | 20000000 | 1000000 | 10150 | 16733 | 16733 | 0 | 1624 | 0 | 15109 |
| D11 | S1 | RPP30-HEX | 11444 | 228883 | 15174 | 9605 | 16773 | 16772 | 1 | 1666 | 0 | 15106 |

**Table S2.** Orthogonal testing data on Ultra-dPCR.

| **Dilution** | **Sample_vol_ul** | **channel1_count** | **channel2_count** | **median_signal_channel1** | **median_noise_channel1** | **median_signal_channel2** | **median_noise_channel2** |
| --- | --- | --- | --- | --- | --- | --- | --- |
| NTC | 52.9956 | 1822 | 3 | 3067 | 368 | 3067 | 563 |
| NTC | 55.5795 | 1684 | 4 | 6810 | 370 | 3200.5 | 584 |
| NTC | 53.7933 | 1910 | 6 | 5711 | 371 | 4209 | 606 |
| NTC | 54.3236 | 1948 | 7 | 5892 | 369 | 7664 | 631 |
| S10 | 52.077 | 1918 | 174 | 6572 | 373 | 8240 | 550 |
| S10 | 54.3553 | 1950 | 170 | 6555 | 375 | 7876.5 | 570 |
| S10 | 52.7465 | 1939 | 149 | 5613 | 370 | 7355.5 | 602 |
| S10 | 54.3407 | 1959 | 143 | 6067 | 372 | 7667 | 612 |
| S9 | 50.4001 | 1863 | 2186 | 6711.5 | 368 | 8172 | 542 |
| S9 | 53.515 | 1968 | 2370 | 6858.5 | 374 | 8505 | 564 |
| S9 | 52.1148 | 1872 | 2203 | 5865.5 | 374 | 7689 | 590 |
| S9 | 53.9695 | 1973 | 2277 | 5865 | 376 | 7824.5 | 598 |
| S8 | 46.9913 | 1655 | 21732 | 6151 | 358 | 7536 | 537 |
| S8 | 53.5456 | 2007 | 25659 | 5910 | 361 | 7629 | 577 |
| S8 | 52.5247 | 1895 | 24678 | 5697 | 373 | 7794.5 | 652 |
| S8 | 53.7508 | 1876 | 25702 | 6146.5 | 366 | 8069 | 611 |
| S7 | 53.4778 | 1899 | 52492 | 6994 | 390 | 8686.5 | 600 |
| S7 | 54.5839 | 2108 | 54689 | 6599 | 394 | 8372 | 636.5 |
| S7 | 52.1288 | 2065 | 53101 | 5783 | 389 | 8046 | 643 |
| S7 | 54.2903 | 1980 | 54221 | 6495 | 392 | 8517.5 | 648 |
| S6 | 52.4052 | 1859 | 118844 | 6626 | 375 | 8574 | 639 |
| S6 | 55.6687 | 1941 | 126902 | 6440 | 387 | 8551 | 690 |
| S6 | 53.8211 | 1861 | 122789 | 6264 | 374 | 8437 | 687 |
| S6 | 53.488 | 1711 | 122521 | 5708 | 376 | 7971 | 684 |
| S5 | 50.9371 | 1774 | 152460 | 6858.5 | 383 | 8613 | 651 |
| S5 | 55.5952 | 2048 | 165562 | 6489 | 384 | 8627 | 688 |
| S5 | 52.6996 | 1965 | 160076 | 7330 | 384 | 10260 | 722 |
| S5 | 53.508 | 1930 | 161063 | 6241 | 385 | 8498 | 708 |
| S4 | 52.3932 | 1661 | 164251 | 6891 | 381 | 9290 | 692 |
| S4 | 54.9414 | 1860 | 171007 | 6741 | 382 | 9184 | 701 |
| S4 | 52.5204 | 1851 | 167586 | 6238 | 383 | 8420 | 719 |
| S4 | 53.9262 | 1939 | 170839 | 5963.5 | 388 | 8243 | 719 |
| S3 | 54.6043 | 1913 | 189074 | 6715.5 | 386 | 8692 | 694 |
| S3 | 52.3681 | 1945 | 181751 | 7261 | 401 | 9216 | 710 |
| S3 | 51.0249 | 1822 | 182480 | 5802 | 390 | 7932 | 721 |
| S3 | 54.4086 | 1881 | 188421 | 6144.5 | 383 | 8566 | 739 |
| S2 | 53.279 | 1900 | 192130 | 6676 | 392 | 8697 | 699 |
| S2 | 53.3843 | 1810 | 191515 | 7375 | 394 | 9574 | 736 |
| S2 | 50.931 | 1875 | 185897 | 5842 | 403 | 7899 | 750 |
| S2 | 53.5131 | 1975 | 191108 | 6293 | 393 | 8881 | 764 |
| S1 | 50.7397 | 1967 | 235454 | 6448 | 396 | 8256 | 710 |
| S1 | 53.458 | 1858 | 244005 | 7020 | 397 | 9363 | 755 |
| S1 | 50.1706 | 1829 | 237441 | 6044 | 407 | 8429 | 822 |
| S1 | 52.7064 | 1784 | 241517 | 5930 | 401 | 8143 | 773 |

